# Circadian rhythmicity and reinforcement processing: a dataset of MRI, fMRI, and behavioral measurements

**DOI:** 10.1101/2025.01.27.635143

**Authors:** Patrycja Scislewska, Michal Rafal Zareba, Julia Lengier, Aaron E. Schirmer, Piotr Bebas, Iwona Szatkowska

**Affiliations:** Department of Animal Physiology, Institute of Experimental Zoology, Faculty of Biology, University of Warsaw, 02-096 Warsaw, Poland; Laboratory of Emotions Neurobiology, Nencki Institute of Experimental Biology, Polish Academy of Sciences, 02-093 Warsaw, Poland; Department of Basic and Clinical Psychology and Psychobiology, Jaume I University, 12-006 Castellon de la Plana, Spain; Department of Biology, Northeastern Illinois University, 5500 N. St. Louis Ave, Chicago, IL 60625, USA

## Abstract

Circadian rhythmicity is a complex phenomenon that influences human behavior, emotionality and brain activity. A detailed description of individual differences in circadian rhythmicity could inform the design of educational systems, shift-worker schedules, and daily routines. Here we present a comprehensive dataset for studying diurnal rhythms and their relationship with human behavior. Thirty seven male participants (aged 20-30) filled in validated psychometric questionnaires assessing characteristics of the circadian rhythm, sleep quality, emotionality, personality traits, reward-punishment processing and attention deficits. Moreover, we acquired high-resolution anatomical T1-weighted images using magnetic resonance imaging (MRI), B0 fieldmaps for distortion corrections, and functional MRI (fMRI) during the Monetary Incentive Delay (MID) task, which is a common paradigm to assess human neural reinforcement processing. All files are organized in Brain Imaging Data Structure (BIDS) and openly available on OpenNeuro.org. The validation of all data confirmed high-quality of described dataset. The various psychological measures combined with neuroimaging data provide strong foundation for exploring emotionality, affective processing, and attention in the context of brain activity and circadian influences.

## Background & Summary

The circadian rhythmicity in humans is a complex phenotype that encompasses multiple dimensions. Full description of the daily rhythm requires a characterization of three distinct components: chronotype, distinctness (fluctuations of the perceived energy level during the day, i.e., amplitude of diurnal variations in mood, motivation, and cognitive functioning), and morning affect (energy and alertness soon after waking, irrespective of wake-up time)^1^. However, the majority of investigations used a unidimensional assessment of morningness-eveningness (chronotypes)^2^. In these studies, eveningness has been associated with negative emotionality, propensity to depression and anxiety, neuroticism, and reduced reward responsiveness^3–6^. Additionally, altered reward-related brain functioning has been observed in people with evening chronotype^7–9^. Neuroticism, as well as propensity to depression and anxiety, are associated with motivation-related deficits, which are reflected in decreased responsiveness to rewarding stimuli, reduced approach-related behaviors, enhanced sensitivity to punishment, and increased avoidance behavior^10^. In addition, previous research revealed that, despite the people with evening chronotype showing higher intelligence^11–13^ and cognitive ability^14^, they have worse academic achievements^15,16^. These findings suggest that incentive motivation is modulated by the morningness-eveningness dimension of the circadian system. Recent research has also suggested the importance of distinctness and morning affect for these processes^17,18^. For example, distinctness negatively correlates with life satisfaction and positively correlates with negative emotionality^5,17,19,20^. Moreover, our recent study revealed that gray matter volume and cortical thickness in the left primary visual cortex are negatively correlated with the distinctness, regardless of chronotype. Furthermore, the association of distinctness with gray matter volume in the right middle temporal gyrus differs between morning- and evening-chronotype groups^21^. As both areas have been implicated in the processing of positive and negative emotional stimuli, the existing evidence points toward the important role of circadian dimensions other than chronotype in reinforcement processing. Nevertheless, such studies remain sparse.

To further explore these aspects of human diurnal rhythmicity, we conducted a study with two main components: examining the subjects’ behavior using psychometric questionnaires and measuring brain activity using functional Magnetic Resonance Imaging (fMRI). The characteristics of the individual circadian rhythmicity were assessed using the Morningness-Eveningness Stability-Scale improved (MESSi) questionnaire, and sleep quality was controlled using the Athens Insomnia Scale. We recruited 37 male participants (aged 20-30) and collected diverse psychometric data. This allowed us to assess individual differences in sensitivity to punishment and reward, personality traits, positive and negative affect, behavioral inhibition system and behavioral activation system (BIS/BAS), anxiety levels, and attention deficits (for details, see: Methods). Additionally, participants also completed an fMRI version of Monetary Incentive Delay task (MID task), which is a widely used paradigm for assessing reward- and punishment-motivated behavior^22,23^.

Although we shared the preliminary results of our analyses as a preprint titled “High distinctness of circadian rhythm is related to negative emotionality and enhanced neural processing of punishment-related information in men” ^24^, the dataset offers significant opportunities for reuse and extension. The broad range of collected psychological data provides a foundation for research focusing on various aspects of human emotionality, affective processing, or even attention disorders.

All of the described data are openly available on OpenNeuro.org, organized according to the standardized Brain Imaging Data Structure (BIDS) format, which facilitates integration with other datasets and opens novel directions for future research projects.

## Methods

### Participants

We invited 37 healthy young men (age 20-30) to participate in the study of circadian-related differences in affective processing and psychological traits. Participants were reached via social media advertisements and local flyers. Subjects were screened before data collection to ensure that all had normal or corrected-to-normal vision, were drug-free, and had no self-reported history of neurological or psychiatric disorders, as well as MRI contraindications. Recruited participants were compensated for their time with a guaranteed payment of 50 PLN (Polish Zloty, 1 PLN equals 0.30 USD). To enhance participants’ motivation to engage in the task, we informed them that the total final payout would depend on their performance in the experiment (up to 110 PLN; 30 USD), and would be received after the experimental session.

The described protocol was approved by the Rector’s Committee for the Ethics of Research Involving Human Participants at the University of Warsaw (decision no. 230/2023) and was performed in accordance with the Declaration of Helsinki. Written informed consent prior to experimentation was obtained from all participants.

### Procedures

#### Overview

The described study consisted of two parts: an online psychometric assessment and an in-person MRI/fMRI scanning session. At the beginning of the study, all participants received individual, 4-digit, random ID codes, which allowed us to combine both psychological and neurobiological data while maintaining participants’ anonymity. All used questionnaires were distributed to the participants using the Psytoolkit online tool^25,26^ (for details, see: Procedures, Behavioral data collection). Participants were asked to complete all surveys before the scanning session. During the appointment, participants underwent a mock MRI training session while performing the MID task (for details, see: Procedures, Monetary Incentive Delay task). This first pre-scanning attempt allowed them to become familiar with the task, to learn the values assigned to specific graphical symbols (cues), and to practice using the response pads (each subject was to respond using the index finger of the right hand). Moreover, during this mock session, we measured the participants’ reaction times to set their individual threshold, ensuring 60% efficiency in the MID task.

Following the training, an actual scanning session was conducted. The entire scanning session lasted approx. 20 minutes. Each session consisted of an anatomical T1-weighted scan (6 min), a fieldmap, and a functional scan during the MID task (13 min comprising 60 trials of 13 seconds each) (for details, see: Procedures, Neuroimaging data collection).

All described procedures were conducted during two weeks in summer, all MRI examinations were carried out from 1 PM to 5 PM.

#### Behavioral data collection

During the study, we collected data on (I) subjective characteristics of the circadian rhythms, using the Polish version of the Morningness-Eveningness-Stability-Scale improved (MESSi) questionnaire^17,27^, (II) sleep quality, using the Athens Insomnia Scale (AIS) questionnaire^28^, (III) individual differences in reinforcement processing, using the Sensitivity to Punishment and Sensitivity to Reward Questionnaire (SPSRQ^29,30^), (IV) personality traits, using NEO-Five Factor Inventory (NEO-FFI^31,32^), (V) behavioral inhibition and activation tendencies, using Behavioral Inhibition System and Behavioral Activation System (BIS/BAS^33,34^), (VI) positive and negative affect, using Positive and Negative Affect Schedule (PANAS^35,36^), (VII) generalized anxiety symptoms, using Generalized Anxiety Disorder (GAD-7^37^) and (VIII) attention deficit and hyperactivity symptoms, using Adult ADHD Self-Report Scale (ASRS^38,39^).

#### Monetary Incentive Delay task

Monetary Incentive Delay task (MID task) is a common method of assessing motivated behavior ^22,23^. During the task performance, participants have to rely on previously learned associations between the cues and monetary outcomes. Therefore, the task is assumed to activate brain regions involved in reward-punishment processing, as well as attention and working memory, such as prefrontal cortex (PFC). The experiment was performed using Presentation software (Version 23.0, Neurobehavioral Systems, Inc., Berkeley, CA, www.neurobs.com).

Each trial in the MID task consists of 5 stages, as shown in Figure 1. At the beginning of each trial, a graphic symbol - cue - with an assigned monetary value is displayed. After the cue, a fixation cross is shown. In the next stage, a screen with a white, filled square appears - this signals that the subject should click the button on the response pad as quickly as possible to get a reward/avoid a punishment (depending on the previously displayed cue). After the reaction (click), the participant gets a feedback screen, displaying the result of a given trial and the total money amount collected so far. After the feedback, another fixation cross appears, and the next trial begins. One trial lasts 13 seconds, and this is repeated for 60 cycles. The anticipatory screen and the reaction screen were defined with 8 pseudo-random combinations of their durations to avoid participants learning the clicking rhythm. The duration of each trial is kept the same by having these combinations always add up to 6 seconds (e.g. 2200 ms + 3800 ms or 2550 ms + 3450 ms), (Figure 1).

**Figure 1.**
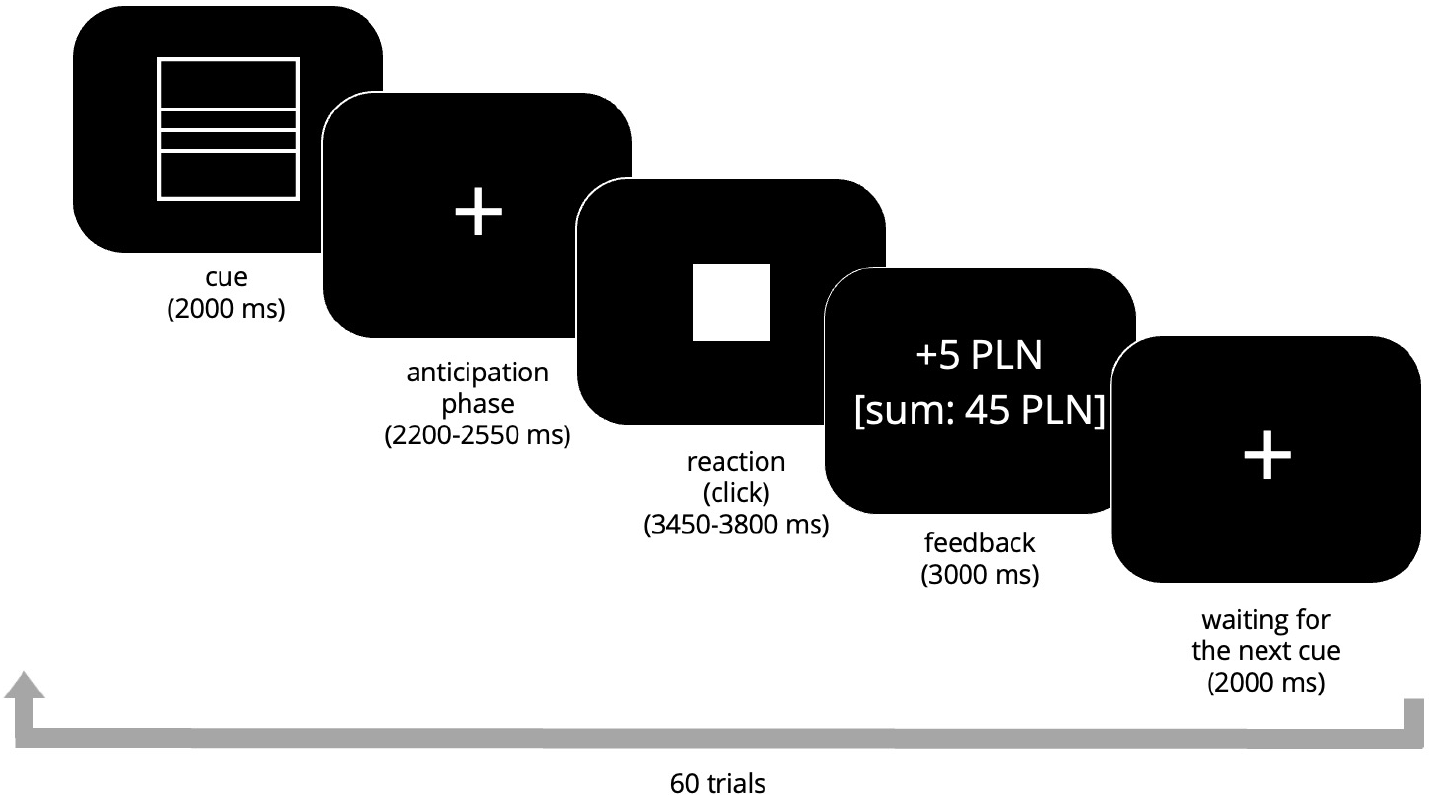
A schematic of Monetary Incentive Delay (MID) task used in the described study.

To study the aspects of neural reinforcement processing (reward and punishment), we designed an experiment consisting of the MID task (for rewards and punishment processing) combined with the oddball paradigm (for keeping participants focused on cue processing). The task consisted of 60 trials: 28 reward trials (square with 3 lines represents big win: +5 PLN, square with 1 line represents a small win: +1 PLN), 28 punishment trials (circle with 3 lines represents big loss: –5 PLN, circle with 1 line represents small loss: –1 PLN), and 4 oddball trials to maintain participants’ focus (triangle, +/-10 PLN) the expected behavior was to inhibit the clicking reaction. The sequence in which the cues were presented in the task was pseudo-randomized but consistent across all subjects. A summary of cues is shown in Figure 2.

**Figure 2.**
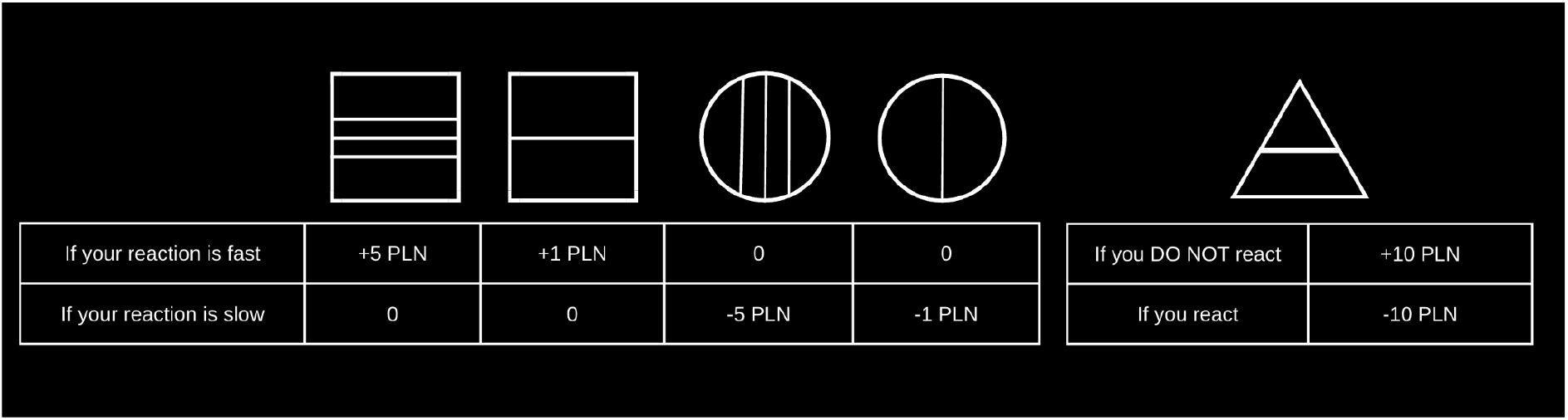
Graphical cues from the Monetary Incentive Delay (MID) task. Participants learned the monetary values of these cues during a training session. At the beginning of the scanning session, participants were shown this screen to refresh the meaning of each cue. However, during the MID task, participants had to rely on their memory.

#### Neuroimaging data collection

The MRI data was acquired using a 3.0 Tesla MAGNETOM Prisma Siemens scanner at the Laboratory of Brain Imaging (LOBI), Neurobiology Center at the Nencki Institute of Experimental Biology Polish Academy of Science. Scans were performed using a 32-channel head coil.

For each subject, we obtained a high-resolution T1-weighted anatomical scan using Gradient Echo with Inversion Recovery sequence (GR\IR). Scans parameters were: slice thickness = 1 mm, echo time (TE) = 0.00296 s, repetition time (TR) = 2.3 s, inversion time (TI) = 0.9 s, isotropic voxel size = 1 ✕ 1 ✕ 1 mm^3^, matrix 208 ✕ 240 with 256 slices. During the performance of the MID task, functional T2*-weighted scans sensitive to blood-oxygenation-level-dependent (BOLD) contrast were collected. The fMRI data were acquired using an echo-planar imaging (EPI) sequence with a multi-band acceleration factor of 2. The scan parameters were as follows: TR = 2.0 s, TE = 0.03 s, flip angle = 70°, isotropic voxel size = 2.5 × 2.5 × 2.5 mm^3^, matrix = 84 × 84, with 60 slices per volume and 397 volumes in total. Moreover, to correct for susceptibility-induced distortions in the fMRI data, we collected a single volume spin-echo EPI B0 fieldmaps. These scans were acquired in anterior-to-posterior (AP) and posterior-to-anterior (PA) phase encoding directions. The scan parameters for both directions were: TE = 0.066 s, TR = 8.0 s, flip angle = 90°, slice thickness = 2.5 mm, spacing between slices = 2.5 mm, isotropic voxel size = 2.5 × 2.5 × 2.5 mm^3^, matrix = 84 × 82, with 60 slices per volume.

Original DICOMs were converted to the NIfTI format using the dcm2niix software package^40^. To maintain participants’ anonymity, scans were defaced using pydeface^41^.

## Data Records

All described data records are publicly available as OpenNeuro Dataset ds005479 (https://openneuro.org/datasets/ds005479/versions/1.0.3)^42^. The dataset includes a README file, a dataset description, a participant file with psychometric data, and neuroimaging data. Neuroimaging files are organized according to the Brain Imaging Data Structure (BIDS)^43^, within folders for each participant (<sub-XX>) and specific subfolders scheme: 1) <anat> (raw, defaced anatomical T1-weighted MRI scans), 2) <func> (raw BOLD data and an event file (.tsv file) listing onset times, durations, and types of stimuli), and 3) <fmap> (raw B0 fieldmaps acquired in two phase encoding directions (AP and PA)). Additionally, each subfolder includes a .json file with technical details about the corresponding NIfTI files.

## Technical Validation

To ensure the technical validation of our behavioral and neuroimaging data, we conducted a series of comprehensive analyses. For details about the psychometric dataset, see: Technical Validation, Behavioral data. Neuroimaging data quality was evaluated using a set of image quality metrics: Contrast-to-noise ratio (CNR), framewise displacement (FD), DVARS, temporal Signal-to-noise ratio (tSNR) (for details, see: Technical Validation, Neuroimaging data). Here, we present the results of these validation procedures across all 37 participants. We note that no data were excluded from the published dataset. Users are encouraged to make their own decisions regarding the exclusion or use of specific subgroups of participants, and the provided metrics may be helpful in making those decisions.

### Behavioral data

The described dataset contains psychometric properties collected from 37 men (aged 20-30). There are no missing values or duplications. In Table 1. we provided descriptive statistics for the dataset, including the mean, standard deviation (SD), minimum and maximum scores obtained by participants, first quartile (Q1, 25%), second (Q2, 50%), and third quartile (Q3, 75%), skewness, kurtosis, and results of the Kolmogorov-Smirnov (KS) test for normality (D-statistics and p-values). According to the KS test, all variables are normally distributed (p-value>0.05). In Figure 3. histograms and corresponding density curves are shown for all variables.

**Table 1.** The descriptive statistics for all psychometric variables from the dataset. Psychometric parameters were assessed using the Morningness-Eveningness-Stability-Scale improved questionnaire(MESSi^17,27^) with subscales of Distinctness, Eveningness, and Morning affect, the Sensitivity to Punishment and Sensitivity to Reward Questionnaire (SPSRQ^29,30^), NEO-Five Factor Inventory (NEO-FFI^31,32^) with subscales of Neuroticism, Extraversion, Openness, Agreeableness, and Conscientiousness, Generalized Anxiety Disorder (GAD-7^37^), Positive and Negative Affect Schedule (PANAS^35,36^) Behavioral Inhibition System and Behavioral Activation System (BIS/BAS^33,34^) with subscales of BAS drive, BAS fun seeking, and BAS reward responsiveness, the Athens Insomnia Scale questionnaire (AIS)^28^, and Adult ADHD Self-Report Scale (ASRS^38,39^). Abbreviations: SD - standard deviation, min - minimum score, max - maximum score, KS - Kolmogorov-Smirnov test.

|  | Count | Mean | SD | min | 25% | 50% | 75% | max | Skewness | Kurtosis | KS statistics (D-statistics) | KS p-value |
| --- | --- | --- | --- | --- | --- | --- | --- | --- | --- | --- | --- | --- |
| **Age** | 37 | 24.432 | 2.089 | 20.0 | 23.0 | 25.0 | 25.0 | 30.0 | 0.208 | 0.530 | 0.165 | 0.236 |
| **Distinctness** | 37 | 18.243 | 3.068 | 10.0 | 17.0 | 18.0 | 20.0 | 25.0 | -0.625 | 0.711 | 0.144 | 0.389 |
| **Eveningness** | 37 | 18.622 | 4.651 | 10.0 | 15.0 | 20.0 | 22.0 | 24.0 | -0.745 | -0.902 | 0.184 | 0.143 |
| **Morning affect** | 37 | 14.27 | 4.953 | 5.0 | 10.0 | 14.0 | 19.0 | 24.0 | 0.203 | -0.942 | 0.116 | 0.656 |
| **Sensitivity to punishment** | 37 | 4.73 | 3.339 | 0.0 | 2.0 | 5.0 | 7.0 | 11.0 | 0.271 | -0.937 | 0.118 | 0.643 |
| **Sensitivity to reward** | 37 | 4.595 | 1.691 | 2.0 | 3.0 | 4.0 | 6.0 | 8.0 | 0.097 | -0.955 | 0.151 | 0.334 |
| **Neuroticism** | 37 | 25.216 | 7.983 | 13.0 | 19.0 | 25.0 | 32.0 | 41.0 | 0.278 | -1.070 | 0.143 | 0.398 |
| **Extraversion** | 37 | 27.027 | 7.481 | 12.0 | 22.0 | 26.0 | 32.0 | 42.0 | 0.099 | -0.569 | 0.072 | 0.984 |
| **Openness** | 37 | 31.351 | 6.672 | 15.0 | 26.0 | 33.0 | 36.0 | 42.0 | -0.502 | -0.522 | 0.138 | 0.441 |
| **Agreeableness** | 37 | 29.351 | 7.166 | 12.0 | 26.0 | 30.0 | 34.0 | 41.0 | -0.801 | 0.066 | 0.131 | 0.511 |
| **Conscientiousness** | 37 | 27.973 | 7.489 | 14.0 | 23.0 | 28.0 | 31.0 | 46.0 | 0.248 | -0.167 | 0.123 | 0.587 |
| **GAD** | 37 | 5.622 | 3.294 | 0.0 | 3.0 | 6.0 | 7.0 | 13.0 | 0.264 | -0.378 | 0.149 | 0.352 |
| **Positive affect** | 37 | 33.027 | 6.768 | 16.0 | 30.0 | 33.0 | 36.0 | 50.0 | -0.203 | 0.880 | 0.148 | 0.359 |
| **Negative affect** | 37 | 19.757 | 5.58 | 11.0 | 16.0 | 18.0 | 25.0 | 30.0 | 0.321 | -0.947 | 0.164 | 0.244 |
| **BAS drive** | 37 | 9.351 | 2.7 | 5.0 | 7.0 | 9.0 | 11.0 | 16.0 | 0.47 | -0.302 | 0.119 | 0.625 |
| **BAS fun** | 37 | 10.595 | 1.691 | 7.0 | 9.0 | 11.0 | 12.0 | 14.0 | -0.323 | -0.524 | 0.175 | 0.182 |
| **BAS reward** | 37 | 14.541 | 2.567 | 9.0 | 13.0 | 15.0 | 16.0 | 19.0 | -0.387 | -0.375 | 0.121 | 0.612 |
| **BIS** | 37 | 19.459 | 4.161 | 8.0 | 17.0 | 19.0 | 23.0 | 27.0 | -0.42 | 0.15 | 0.108 | 0.744 |
| **AIS** | 37 | 5.595 | 3.166 | 1.0 | 3.0 | 5.0 | 7.0 | 16.0 | 1.092 | 1.578 | 0.125 | 0.565 |
| **ASRS** | 37 | 34.378 | 10.505 | 18.0 | 26.0 | 32.0 | 41.0 | 61.0 | 0.473 | -0.423 | 0.112 | 0.703 |

**Figure 3.**
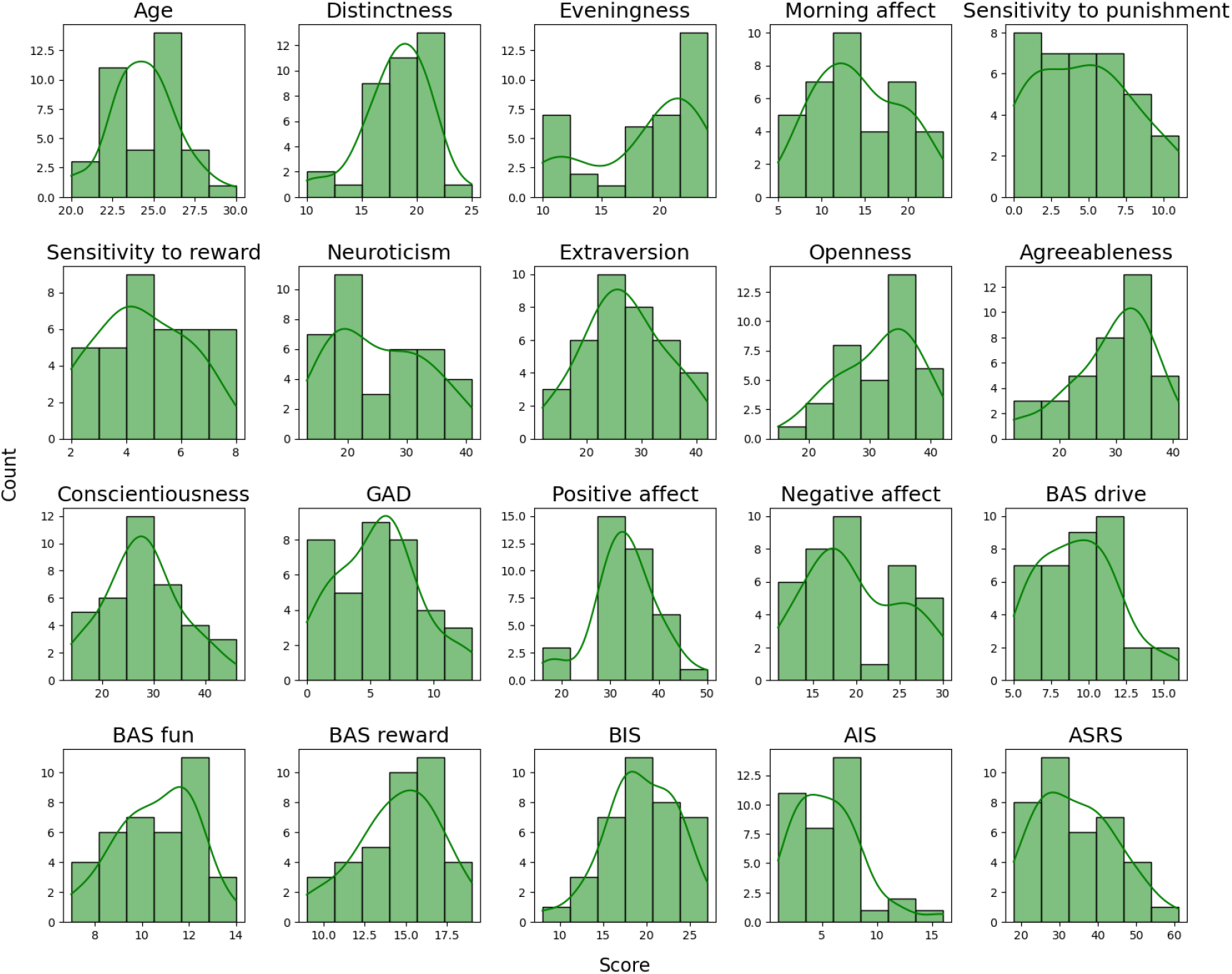
Histograms and density curves for all variables in the described dataset. For ease of comparison, each scale was divided into 6 bins. Psychometric parameters were assessed using the Morningness-Eveningness-Stability-Scale improved questionnaire(MESSi^17,27^) with subscales of Distinctness, Eveningness, and Morning affect, the Sensitivity to Punishment and Sensitivity to Reward Questionnaire (SPSRQ^29,30^), NEO-Five Factor Inventory (NEO-FFI^31,32^) with subscales of Neuroticism, Extraversion, Openness, Agreeableness, and Conscientiousness, Generalized Anxiety Disorder (GAD-7^37^), Positive and Negative Affect Schedule (PANAS^35,36^) Behavioral Inhibition System and Behavioral Activation System (BIS/BAS^33,34^) with subscales of BAS drive, BAS fun seeking, and BAS reward responsiveness, the Athens Insomnia Scale questionnaire (AIS)^28^, and Adult ADHD Self-Report Scale (ASRS^38,39^).

### Neuroimaging data

The dataset includes raw MRI data, however, to validate the quality of MRI data, we calculated all necessary metrics using preprocessed images. All procedures were performed using FSL tools and custom scripts, as described below. Users are encouraged to apply their own preprocessing steps to tailor the images to their specific analytical approaches.

#### Fieldmap processing

Spin Echo Field Maps were collected with reversed phase-encode blips, resulting in pairs of images with distortions going in opposite directions: anterior-to-posterior (AP) and posterior-to-anterior (PA) phase encoding. From these pairs, the susceptibility-induced off-resonance field was estimated using FSL^44^, the two images were combined into a single corrected one (fslmerge and topup tools).

#### Anatomical data processing

T1-weighted anatomical scans were preprocessed using the fsl_anat pipeline in FSL. Original images were reoriented to match the standard MNI152 orientation using fslreorient2std. Non-brain regions were automatically cropped from the image using robustfov. Intensity inhomogeneities caused by RF/B1 field non-uniformity were corrected using FAST. The T1-weighted images were registered to the MNI152 standard brain template (both linear registration using FLIRT, and non-linear registration using FNIRT). The brain was extracted using a BET tool. The image was segmented into gray matter (GM), white matter (WM), and cerebrospinal fluid (CSF) compartments using FAST.

The quality of images was evaluated by calculating the Contrast-to-Noise ratio (CNR). All participants exhibited high CNR scores (Figure 4.a) - the contrast between gray and white matter is approximately 4.4 times greater than the background noise, confirming that the quality of described images is sufficient to reliably distinguish between GM and WM.

**Figure 4.**
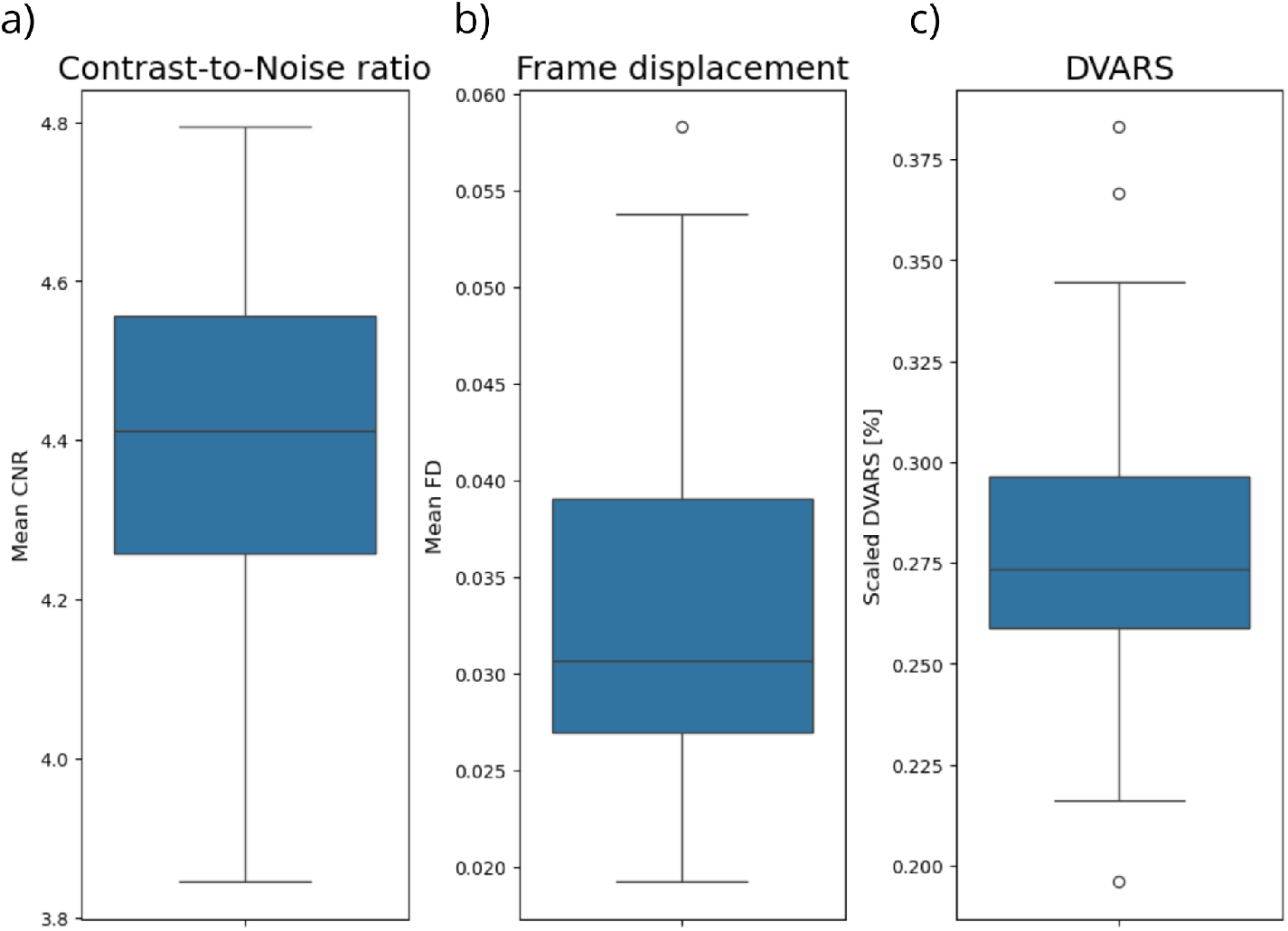
MRI data quality metrics. a) Anatomical data quality was assessed using Contrast-to-Noise ratio (CNR). Obtained CNR scores suggest that quality of T1-weighted images is sufficient for distinguishing tissues and suitable for structural analysis. b) Mean framewise displacement (FD) - metric that detects motion artefacts in functional data. Low values (mostly <0.05 mm) indicate minimal head movement. c) Mean DVARS - metric that evaluates signal stability over time in functional data. Here we present the average percentage change in signal intensity between consecutive time points, normalized to the mean signal intensity. Low values indicate minimal signal fluctuations, which suggests high-quality, low-motion data.

#### Functional data processing

Preprocessing of the functional images included motion correction (MCFLIRT tool), intensity normalization, registration to the MNI152 standard space, and B0 inhomogeneity correction, all performed in FEAT FSL. To preserve image details, no spatial smoothing (FWHM) was applied.

The quality of functional images was assessed using three types of metrics: framewise displacement (FD) and DVARS to detect motion artifacts, temporal Signal-to-Noise ratio (tSNR) to evaluate the stability of the BOLD signal over time. We began examining the functional images quality using FD, which tracks head movement, and DVARS, which tracks changes in image intensity relative to the previous volume(s)^45,46^. Both metrics were calculated using fsl_motion_outliers tool, applying the default outlier threshold of the 75th percentile + 1.5 times the interquartile range (IQR). For better interpretability, we scaled DVARS, by dividing the raw DVARS by mean signal intensity and multiplying by 100%.

There is no universally accepted threshold for movement, but FD thresholds between 0.2 and 0.5 mm are commonly used^46,47^. In the described dataset, both FD and DVARS values were low - FD was mostly < 0.05 mm, scaled DVARS < 0.3%, which indicates minimal head motion and high quality of data (Figure 4.b, 4.c).

For calculating the temporal Signal-to-Noise ratio (tSNR), a metric for signal stability, we used both B0 field-corrected and uncorrected images. Individual and group-level tSNR maps were generated by dividing the mean of a time series by its standard deviation using fslmaths and aligned to the MNI152 standard template using FSL’s FLIRT. Higher tSNR values reflect better stability of the BOLD signal over time. For visualization, a threshold of tSNR > 30 was applied to the mean tSNR map to present regions with the highest signal stability (Figure 5.). We observed that tSNR was high and relatively uniform across the whole brain.

**Figure 5.**
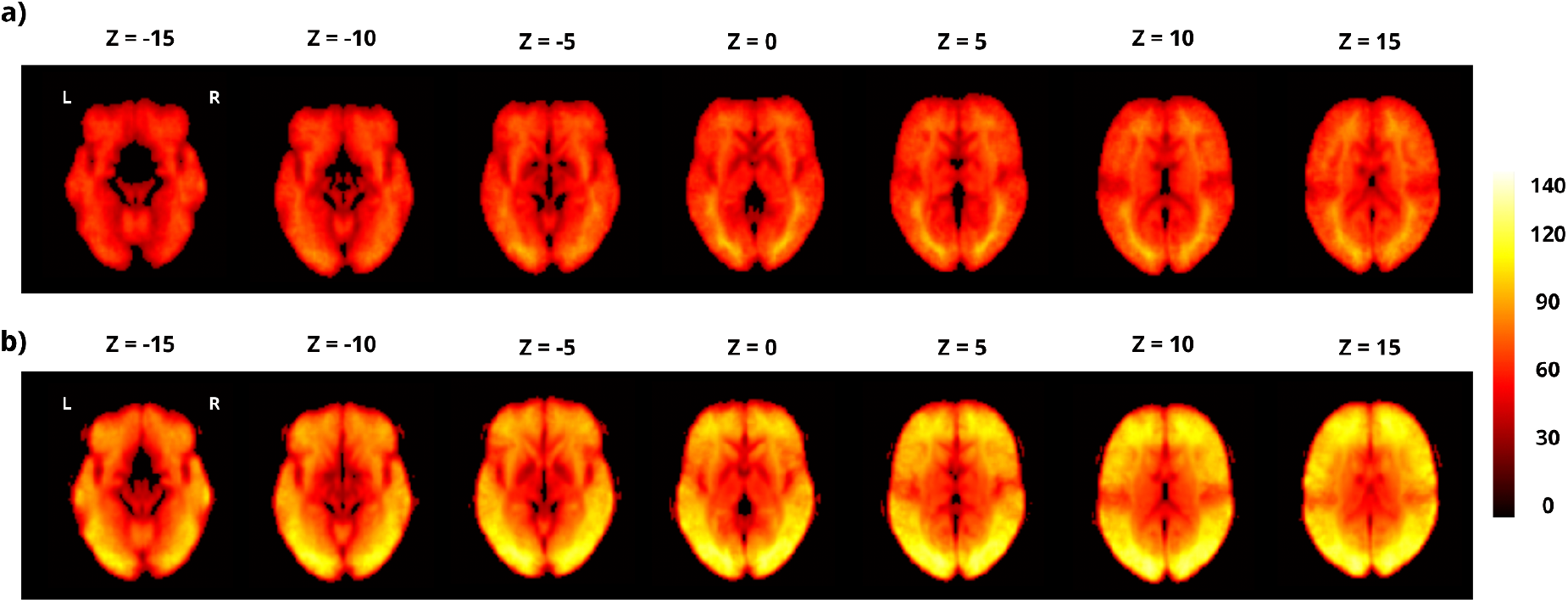
Temporal Signal-to-Noise ratio (tSNR) maps. High values of tSNR indicate a high-quality signal. The tSNR is relatively uniform across the whole brain. a) Map prepared using functional data without distortion correction. b) Map prepared using functional data with B0 inhomogeneity correction applied.

It is worth mentioning here that functional images after or B0 inhomogeneities exhibited higher tSNR values compared to those without distortion correction (Figure 6.), which underlines the importance of using fieldmaps (provided in the described dataset) to improve the quality of signals.

**Figure 6.**
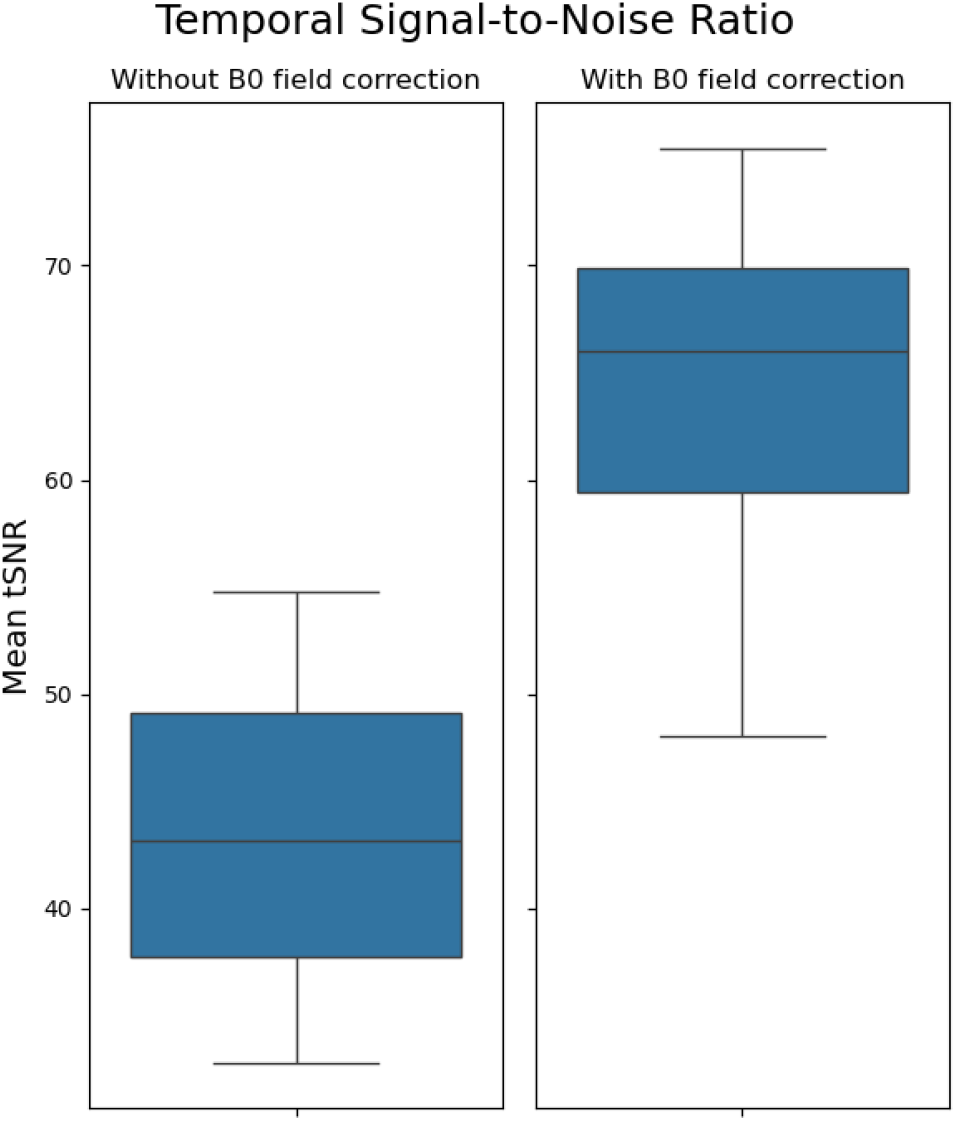
Mean tSNR. Values for functional data with B0 field correction applied are notably higher in comparison with functional data without this correction. This underscores the importance of using the fieldmaps as a crucial step in preprocessing.

## Usage Notes

The described dataset consists of psychological data (on circadian rhythmicity, sleep quality, personality traits, affective processing and emotion processing) and neuroimaging data obtained during performing Monetary Incentive Delay (MID) task from 37 young male participants. All data are organized in the BIDS structure. Data were collected in order to investigate the intriguing relationship between individual circadian rhythms and affective processing. Preliminary results of our analyses are available as a preprint titled “High distinctness of circadian rhythm is related to negative emotionality and enhanced neural processing of punishment-related information in men” ^24^. The paper is currently under peer review. However, the dataset includes a broad range of psychological traits, making it potentially useful for research in other focus areas. No data were excluded from the published dataset, so users can make their own determinations about exclusions.

Data are openly available at OpenNeuro: https://openneuro.org/datasets/ds005479/versions/1.0.3

## Code Availability

To ensure reproducibility and transparency, the code used to perform the technical validation is available on Github: https://github.com/PatrycjaScislewska/Circadian_MRI_dataset. The repository contains custom Python notebooks and Bash scripts to run the open-source FSL tools (fsl_motion_outliers, fslmaths, fslmeants, fslstats).

Acknowledgments

We would like to express gratitude to all the participants of our study. Special thanks go to Dawid Drozdziel, Marta Rodziewicz, and Bartosz Kossowski, PhD, from the Laboratory of Brain Imaging, Nencki Institute of Experimental Biology, Polish Academy of Sciences for their invaluable technical assistance with the MRI data collection.

## Author Contributions

**PS**: Conceptualization, Study design, Methodology, Funding acquisition, Participant recruitment, Data collection, Formal analysis, Writing - Original Draft, Visualization, Figures preparation, **MRZ**: Conceptualization, Methodology, Writing - review & editing **JL**: Participant recruitment, Data collection, Writing - review & editing **AES**: Writing - review & editing, Supervision, **PB**: Writing - review & editing, Supervision, **IS**: Conceptualization, Study design, Writing - review & editing, Supervision.

